# Genetic variants of human platelet antigens in the Indian population from 1029 whole genomes

**DOI:** 10.1101/2022.10.29.514338

**Authors:** Mercy Rophina, Rahul C Bhoyar, Mohamed Imran, Vigneshwar Senthivel, Mohit Kumar Divakar, Anushree Mishra, Bani Jolly, Sridhar Sivasubbu, Vinod Scaria

**Affiliations:** CSIR Institute of Genomics and Integrative Biology (CSIR-IGIB), Mathura Road, Delhi 110025, INDIA; Academy of Scientific and Innovative Research (AcSIR), CSIR-HRDC Campus, Sector 19, Kamla Nehru Nagar, Ghaziabad, Uttar Pradesh 201002, INDIA

**Keywords:** Human Platelet Antigens, Indian population, allele frequency, genotype prevalence

## Abstract

**Background:** Genetic variants in human platelet antigens (HPAs) considered as allo- or auto antigens are associated with various disorders including neonatal alloimmune thrombocytopenia, platelet transfusion refractoriness and post-transfusion purpura. While global differences in genotype frequencies were observed, the distribution of HPA variants in the Indian population are largely unknown. This study aims to explore the landscape of HPA variants in India to provide a basis for risk assessment and management of related complications.

**Materials and methods:** Population specific frequencies of genetic variants associated with the 35 classes of HPAs (HPA-1 to HPA-35) were estimated by systematically analyzing genomic variations of 1029 healthy Indian individuals as well as from global population genome datasets..

**Results:** Allele frequencies of the most clinically relevant HPA systems in the Indian population were found as follows, HPA-1a – 0.89, HPA-1b – 0.15, HPA-2a – 0.94, HPA-2b – 0.05, HPA-3a – 0.66, HPA-3b – 0.36, HPA-4a – 1.00, HPA-4b – 0, HPA-5a – 0.92, HPA-5b – 0.08, HPA-6a – 1.00, HPA-6b – 0, HPA-15a – 0.58 and HPA-15b – 0.42. In addition, HPA-4b allele frequencies were found to be significantly higher in India in comparison to global populations.

**Conclusion:** This study provides the first comprehensive analysis of HPA allele and genotype frequencies using large scale representative whole genome sequencing data of the Indian population.

## Introduction

Platelets are small, anucleate, cells that have a key role in haemostasis. Recent studies have explored their functional roles in inflammation (Leslie 2010), innate and adaptive immunity (Pacheco et al. 2011) and key contributions to a range of disorders including cancer (Leslie 2010). Platelets function through ligand-receptor interactions involving cell surface glycoproteins. Genetic variations occurring in these glycoprotein genes collectively give rise to human platelet antigens (HPAs). HPAs can be recognized as allo- or auto antigens and lead to a range of immune platelet disorders like multitansfusion platelet refractoriness (MPR), fetal and neonatal alloimmune thrombocytopenia (FNAIT) and post transfusion purpura (PTP) (Bianchi et al. 2012), (Pacheco et al. 2011).

HPAs were first described as early as 1960s and about 41 antigens encoded by 8 different platelet glycoproteins namely GPIIIa, GPIbɑ, GPIIIb, GPIa, GPIIb, GPIbβ, CD109 and GPIX have been documented and approved by the International Platelet Immunology Nomenclature Committee of the International Society of Blood Transfusion (ISBT). This includes a total of 6 biallelic systems (HPA-1, HPA-2, HPA-3, HPA-4, HPA-5 and HPA-15) and 29 monoallelic systems (HPA – 6, 7, 8, 9, 10, 11, 12, 13, 14, 16, 17, 18, 19, 20, 21, 22, 23, 24, 25, 26, 27, 28, 29, 30, 31, 32, 33, 34 and 35) (Curtis and McFarland 2014). A vast majority of HPAs are expressed on GPAIIIa, making it the most immunogenic of the platelet glycoproteins (Rozman 2002).

In recent years, with the advent of wider genomic analysis, a large diversity of HPA genotypes have come out of genomic analysis of human ethnic groups from a variety of geographical locations (Silva-Malta et al. 2018),(Jovanovic Srzentic et al. 2021),(Yan et al. 2006). Many of these studies have suggested significant differences in the frequencies between populations. For example, HPA-4b antigen was found to be very rare in Caucasians but more frequent in Asian populations. Anti-HPA-4a and anti-HPA-4b are common and implicated in alloimmune platelet disorders (Jovanovic Srzentic et al. 2021), (Yan et al. 2006), (Tan, Lian, and Nadarajan 2012). The recent availability of genomic data for Indian populations motivated us to analyze the allele and genotype frequencies of HPAs towards understanding frequent and rare alleles in the population .

## Materials and methods

### Datasets used in the study

A comprehensive catalogue of the human platelet antigens approved by the International Platelet Immunology Nomenclature Committee of the International Society of Blood Transfusion (ISBT) was retrieved from Human Platelet Antigen Database. **Table 1** The dataset provides the genomic coordinates and HGVS nomenclature of these reference variants. Genomic variations compiled from whole genome sequences of 1029 self declared healthy individuals from across India were used in the analysis (Jain et al. 2021). This dataset encompasses a total of 55,898,122 variants. Variants associated with HPAs were parsed in this dataset to systematically retrieve the genotype and allele frequencies using bespoke scripts.

**Table 1.**
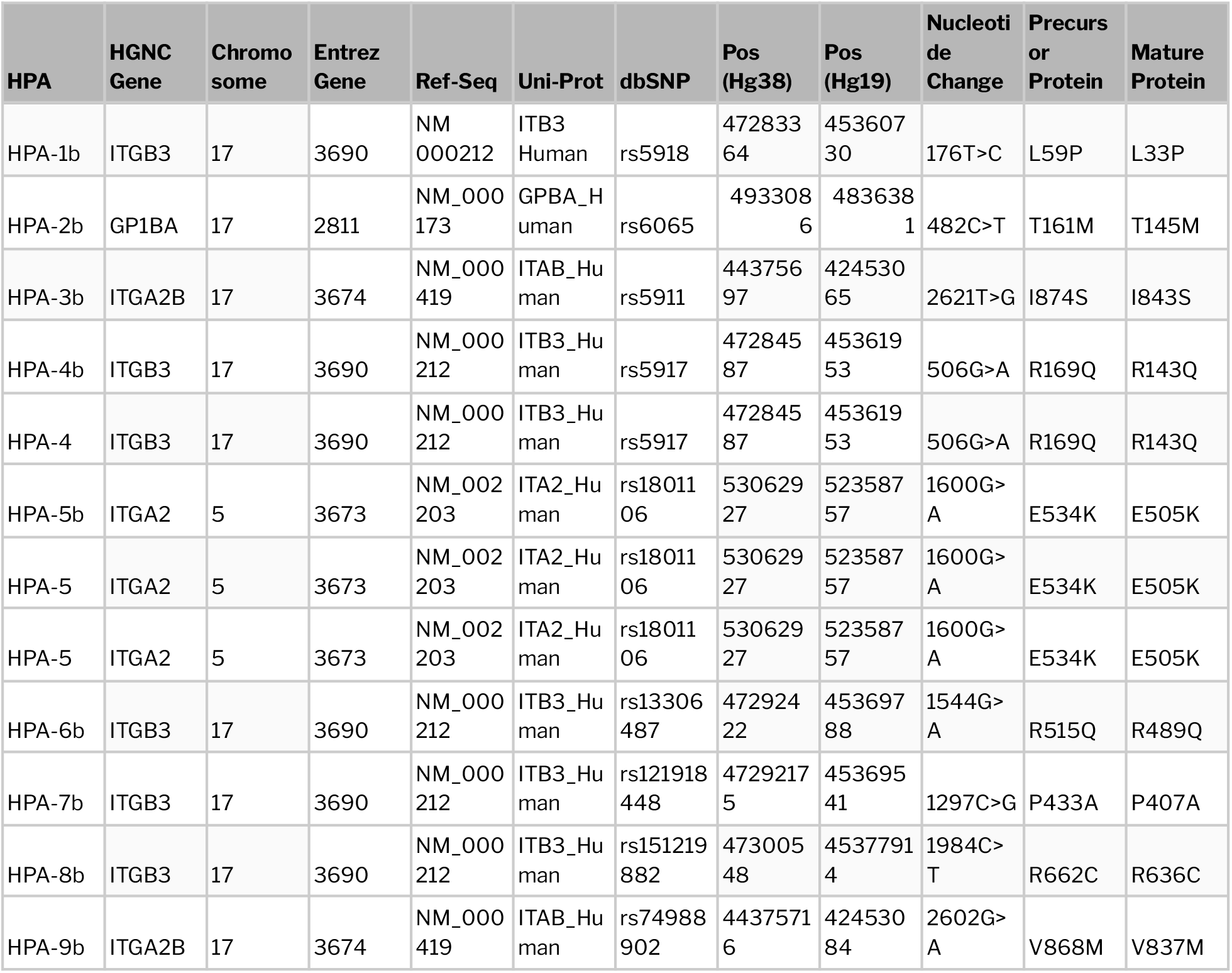
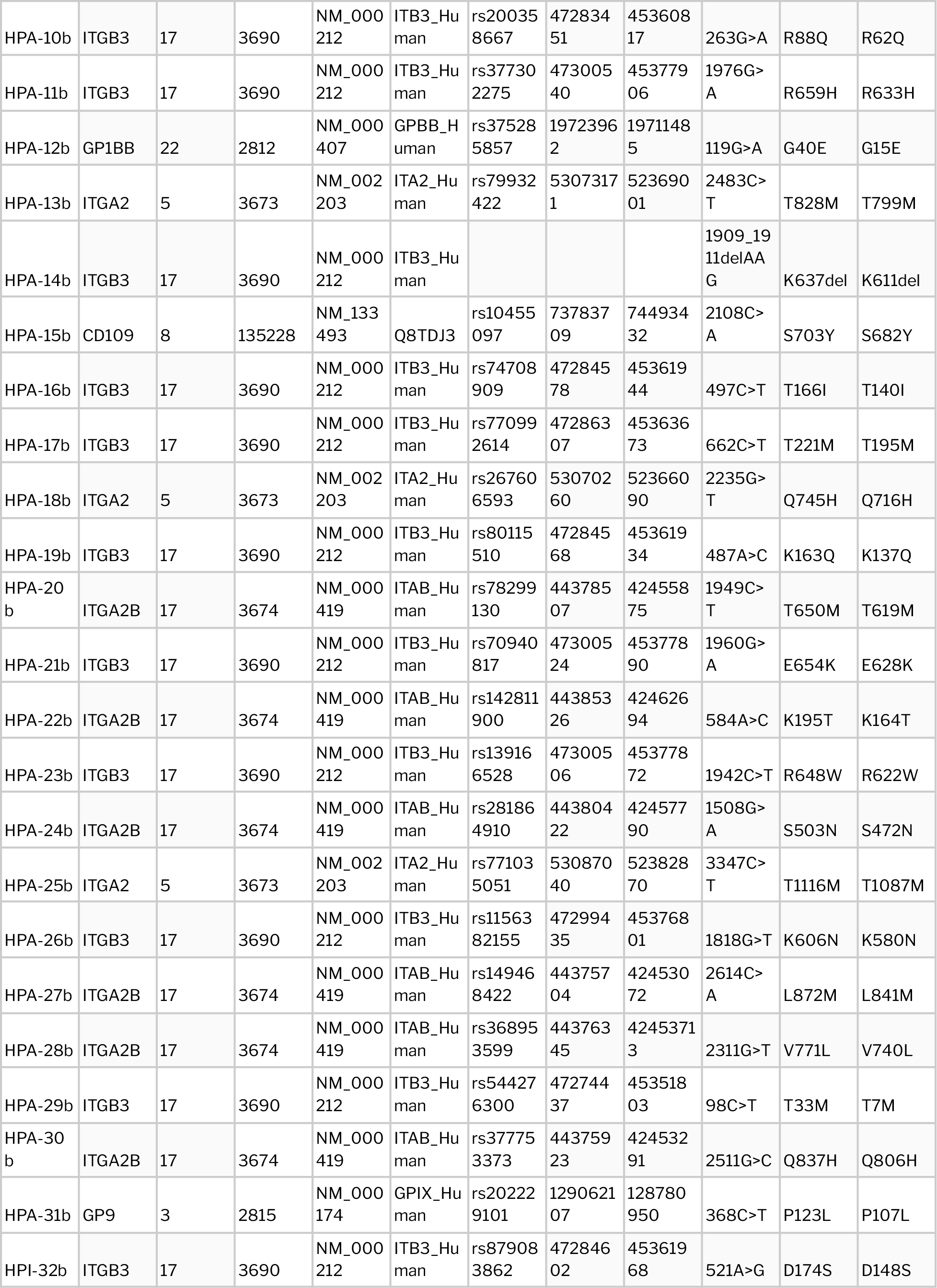
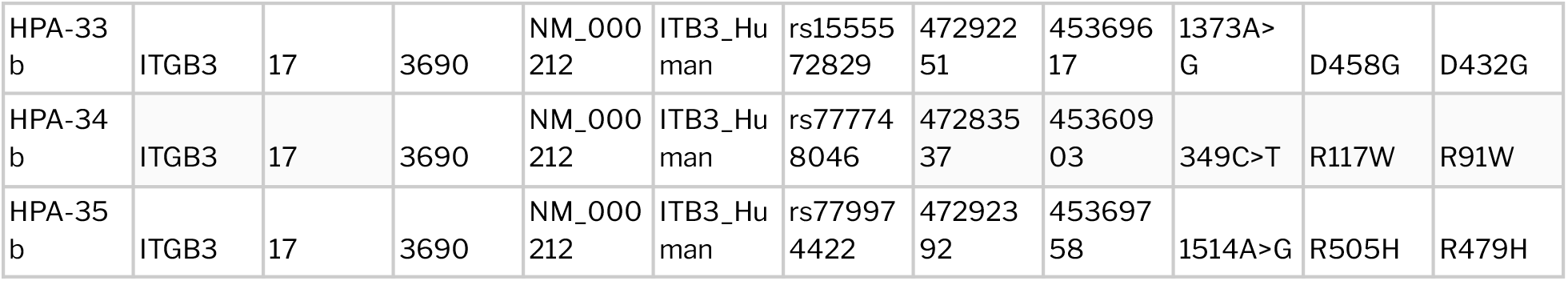
Precise summary of the list of approved HPA variants analysed in the study

### Estimation and comparison of HPA frequencies

Frequencies and zygosity information of HPA variants were retrieved from the dataset of genetic variants compiled from the IndiGenomes resource (Jain et al. 2021). Expected number of genotypes were computed based on Hardy-Weinberg equilibrium calculations. Any deviation of the observed number of genotypes from the expected numbers was evaluated by Chi-square test (p-value < 0.05).

## Results

Frequencies of HPA variants were analysed in a total of 1029 whole genome sequences. **Table 2** summarizes the observed frequencies of HPA variants along with the counts of samples in homozygous and heterozygous states. In the HPA-1 system, HPA-1a/a was observed as the most frequent genotype (79.04%) followed by HPA-1a/b (18.62%) and HPA-1b/b (2.34%). Similarly genotype frequencies observed in classes 2-5 and 15 are as follows, HPA-2a/a – 88.49%, HPA-2a/b – 11.22%, HPA-2b/b – 0.29%, HPA-3a/a – 43.63%, HPA-3a/b – 43.33%, HPA-3b/b – 13.04%, HPA-4a/a – 99.80%, HPA-4a/b – 0.20%, HPA-4b/b – 0%, HPA-5a/a – 85.13%, HPA-5a/b – 14.29%, HPA-5b/b-0.59%, HPA-15a/a – 33.79%, HPA-15a/b – 48.83%, HPA-15b/b – 17.38%. Expected genotype frequencies were duly estimated for HPA-1, 2, 3, 4, 5 and 15 alleles based on Hardy-Weinberg equilibrium. Chi-square test was performed to evaluate significant differences between observed and expected frequencies. Distribution of HPA genotype frequencies in the Indian population tested for Hardy-Weinberg equilibrium is tabulated in **Table 3**. In accordance with the observed p-values, significant deviations of genotypes were seen in HPA-1 (p-val: 0.0094) and −4 (p-val: 0.0001) systems. **Supplementary Tables 1 and 2** elaborate the frequencies of HPA variants reported from various subpopulations of gnomAD (Karczewski et al. 2020) and 1000 genomes project (1000 Genomes Project Consortium et al. 2015). Distribution of HPA allele frequencies among various global populations is graphically depicted in **Figure 1**.

**Table 2.**
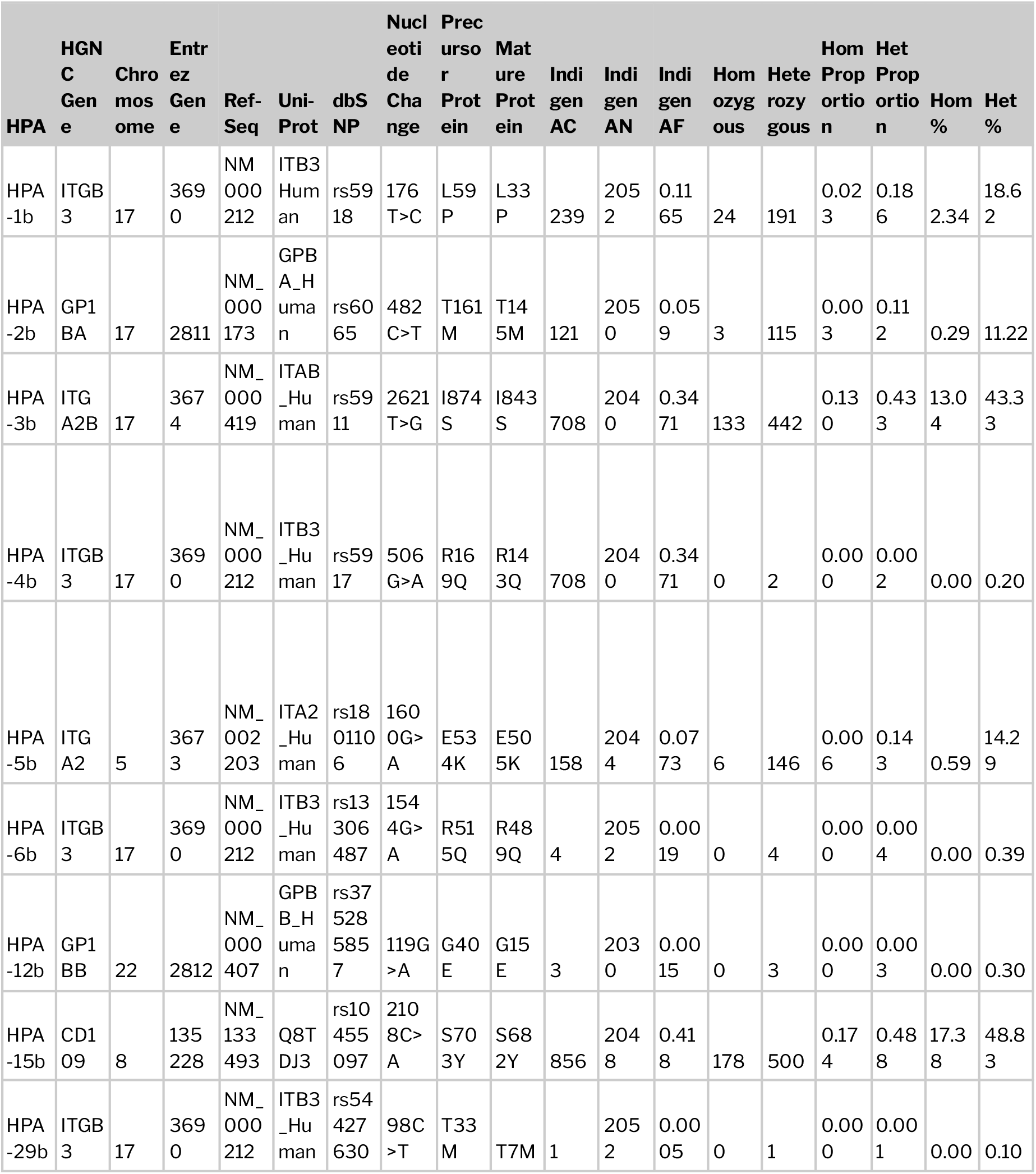
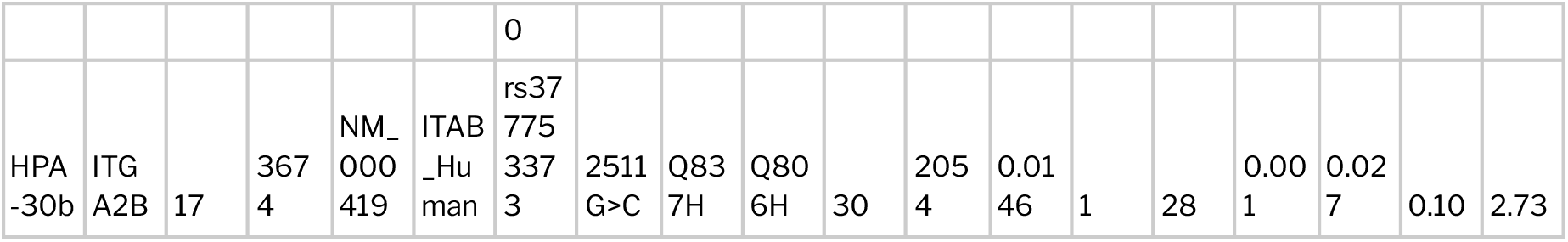
Tabulation of observed frequencies of HPA variants in the Indian population. For HPA systems that are not listed in the table no variants were found in the population.

**Table 3.**
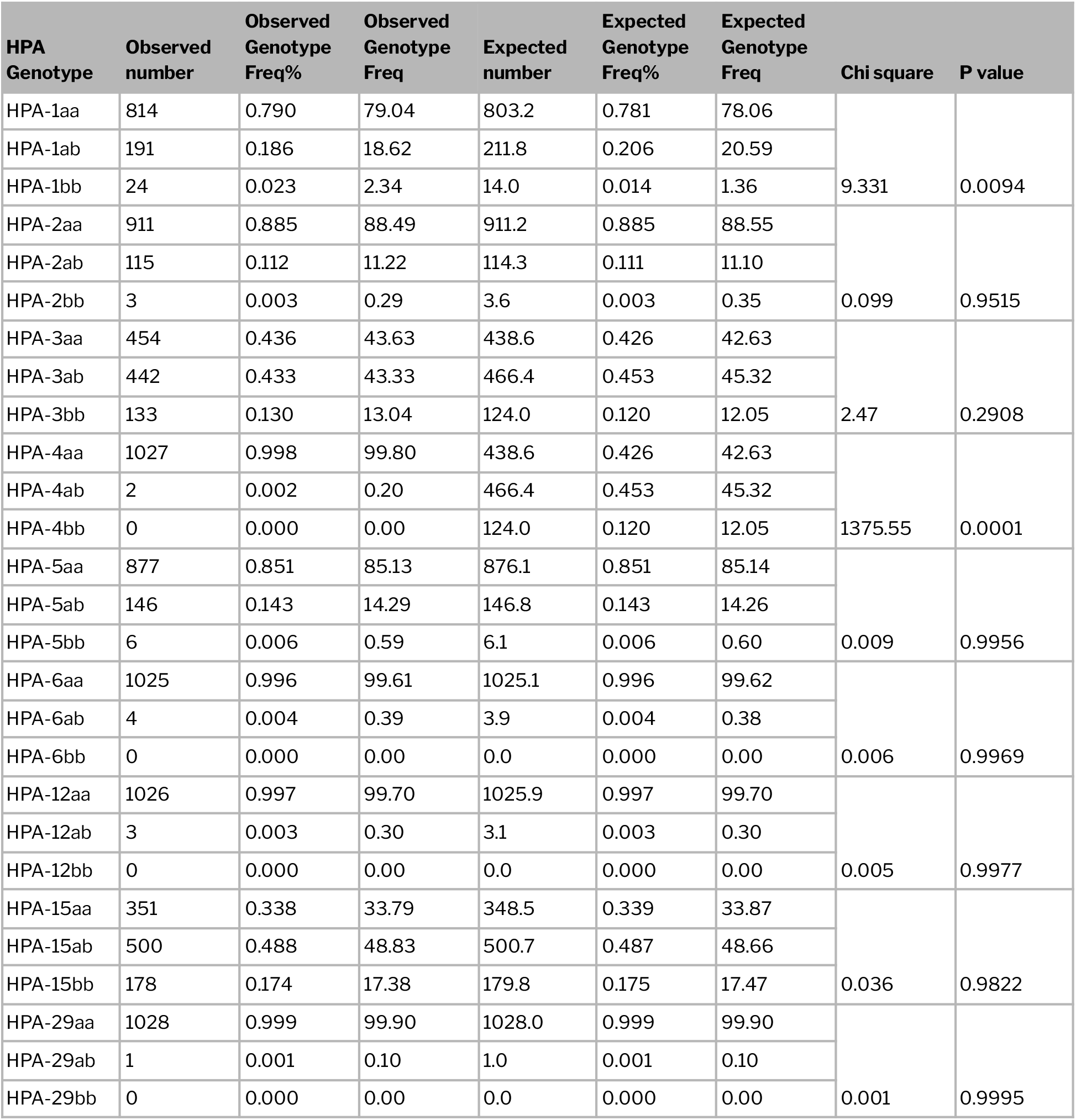
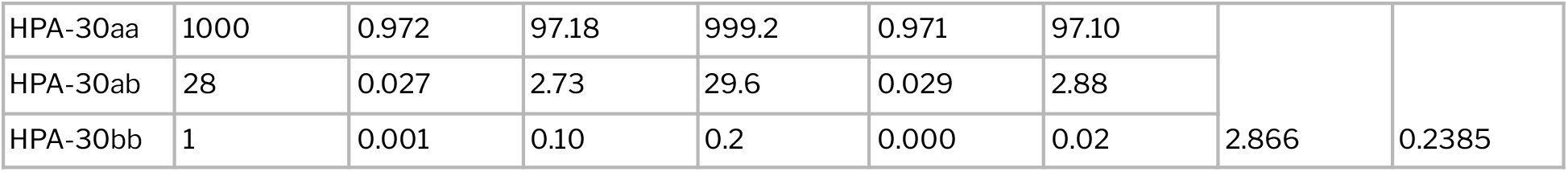
Estimation of Chi squared statistic and P-values using the observed and expected genotype frequencies of various classes of HPA observed in the Indian population

| HPA Genotype | Observed number | Observed Genotype Freq% | Observed Genotype Freq | Expected number | Expected Genotype Freq% | Expected Genotype Freq | Chi square | P value |
| --- | --- | --- | --- | --- | --- | --- | --- | --- |
| HPA-1aa | 814 | 0.790 | 79.04 | 803.2 | 0.781 | 78.06 | 9.331 | 0.0094 |
| HPA-1ab | 191 | 0.186 | 18.62 | 211.8 | 0.206 | 20.59 |  |  |
| HPA-1bb | 24 | 0.023 | 2.34 | 14.0 | 0.014 | 1.36 |  |  |
| HPA-2aa | 911 | 0.885 | 88.49 | 911.2 | 0.885 | 88.55 | 0.099 | 0.9515 |
| HPA-2ab | 115 | 0.112 | 11.22 | 114.3 | 0.111 | 11.10 |  |  |
| HPA-2bb | 3 | 0.003 | 0.29 | 3.6 | 0.003 | 0.35 |  |  |
| HPA-3aa | 454 | 0.436 | 43.63 | 438.6 | 0.426 | 42.63 | 2.47 | 0.2908 |
| HPA-3ab | 442 | 0.433 | 43.33 | 466.4 | 0.453 | 45.32 |  |  |
| HPA-3bb | 133 | 0.130 | 13.04 | 124.0 | 0.120 | 12.05 |  |  |
| HPA-4aa | 1027 | 0.998 | 99.80 | 438.6 | 0.426 | 42.63 | 1375.55 | 0.0001 |
| HPA-4ab | 2 | 0.002 | 0.20 | 466.4 | 0.453 | 45.32 |  |  |
| HPA-4bb | 0 | 0.000 | 0.00 | 124.0 | 0.120 | 12.05 |  |  |
| HPA-5aa | 877 | 0.851 | 85.13 | 876.1 | 0.851 | 85.14 | 0.009 | 0.9956 |
| HPA-5ab | 146 | 0.143 | 14.29 | 146.8 | 0.143 | 14.26 |  |  |
| HPA-5bb | 6 | 0.006 | 0.59 | 6.1 | 0.006 | 0.60 |  |  |
| HPA-6aa | 1025 | 0.996 | 99.61 | 1025.1 | 0.996 | 99.62 | 0.006 | 0.9969 |
| HPA-6ab | 4 | 0.004 | 0.39 | 3.9 | 0.004 | 0.38 |  |  |
| HPA-6bb | 0 | 0.000 | 0.00 | 0.0 | 0.000 | 0.00 |  |  |
| HPA-12aa | 1026 | 0.997 | 99.70 | 1025.9 | 0.997 | 99.70 | 0.005 | 0.9977 |
| HPA-12ab | 3 | 0.003 | 0.30 | 3.1 | 0.003 | 0.30 |  |  |
| HPA-12bb | 0 | 0.000 | 0.00 | 0.0 | 0.000 | 0.00 |  |  |
| HPA-15aa | 351 | 0.338 | 33.79 | 348.5 | 0.339 | 33.87 | 0.036 | 0.9822 |
| HPA-15ab | 500 | 0.488 | 48.83 | 500.7 | 0.487 | 48.66 |  |  |
| HPA-15bb | 178 | 0.174 | 17.38 | 179.8 | 0.175 | 17.47 |  |  |
| HPA-29aa | 1028 | 0.999 | 99.90 | 1028.0 | 0.999 | 99.90 | 0.001 | 0.9995 |
| HPA-29ab | 1 | 0.001 | 0.10 | 1.0 | 0.001 | 0.10 |  |  |
| HPA-29bb | 0 | 0.000 | 0.00 | 0.0 | 0.000 | 0.00 |  |  |
| HPA-30aa | 1000 | 0.972 | 97.18 | 999.2 | 0.971 | 97.10 | 2.866 | 0.2385 |
| HPA-30ab | 28 | 0.027 | 2.73 | 29.6 | 0.029 | 2.88 |  |  |
| HPA-30bb | 1 | 0.001 | 0.10 | 0.2 | 0.000 | 0.02 |  |  |

**Table 4.**
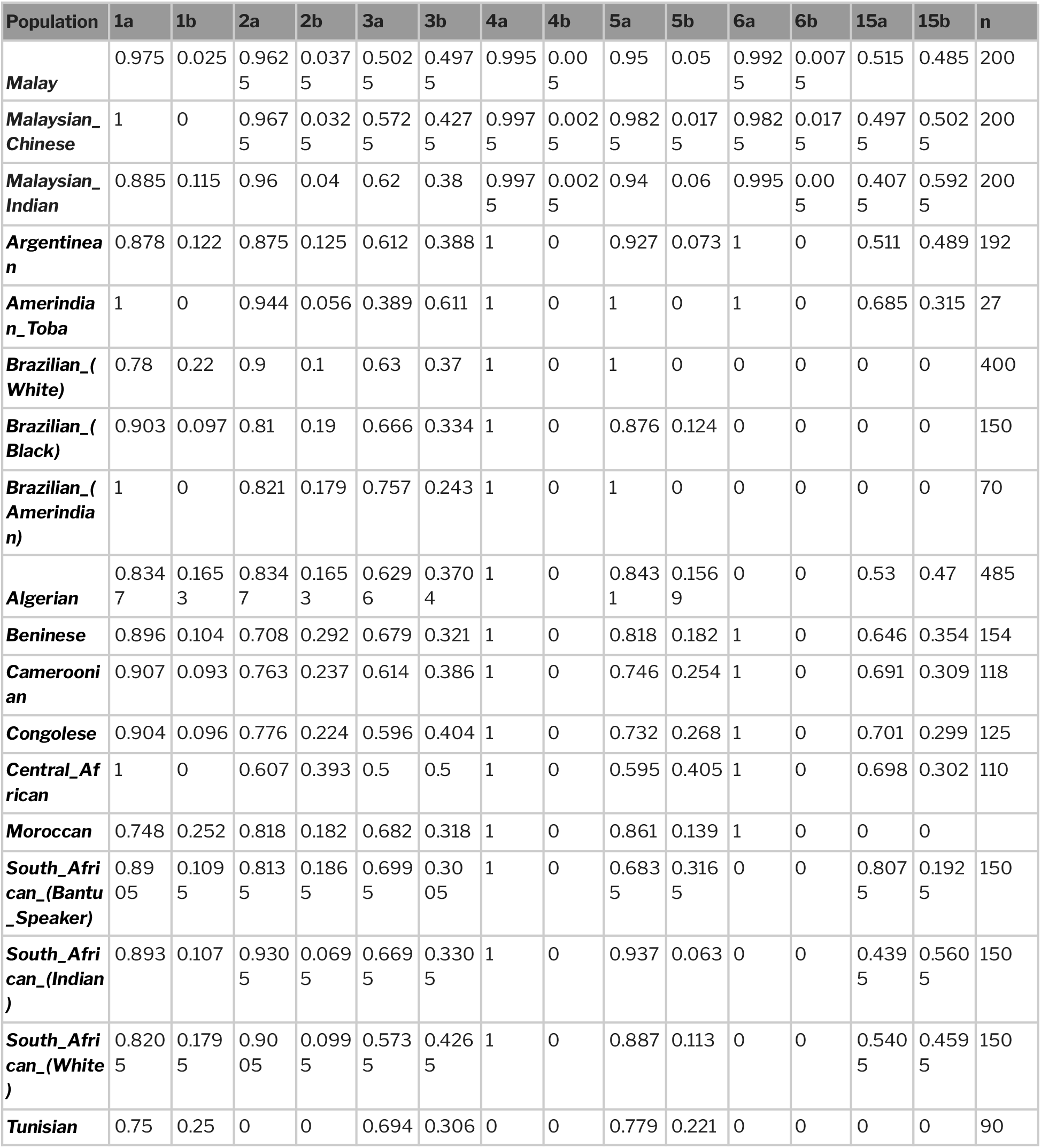
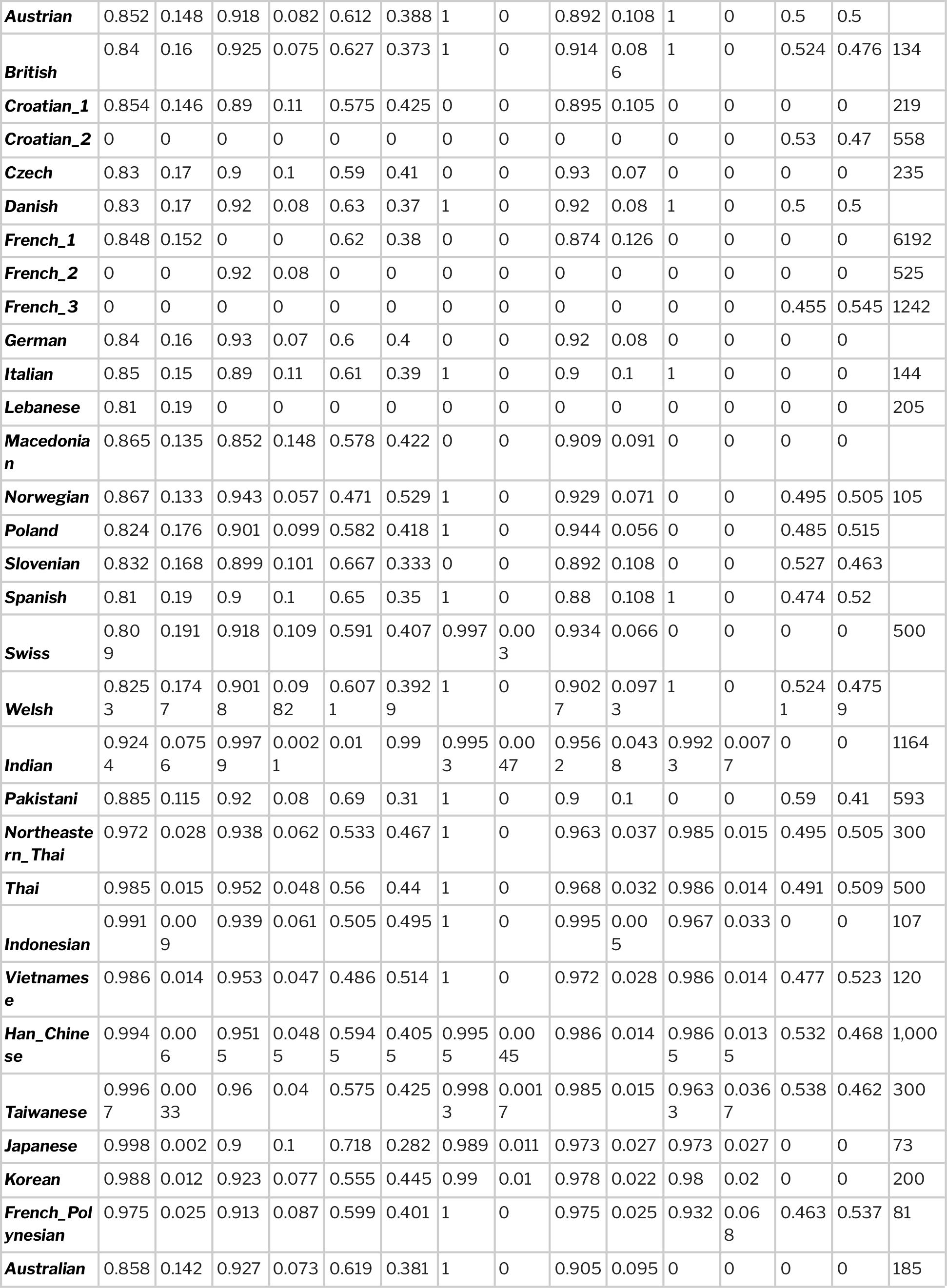
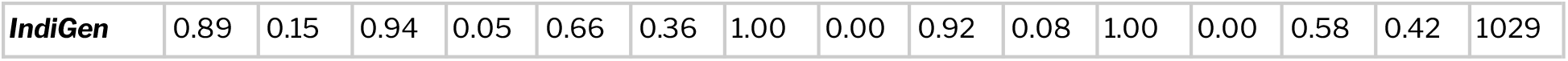
Precise tabulation of allele frequencies of HPA-1,2,3,4,5 and 15 systems across multiple global populations.

**Figure 1.**
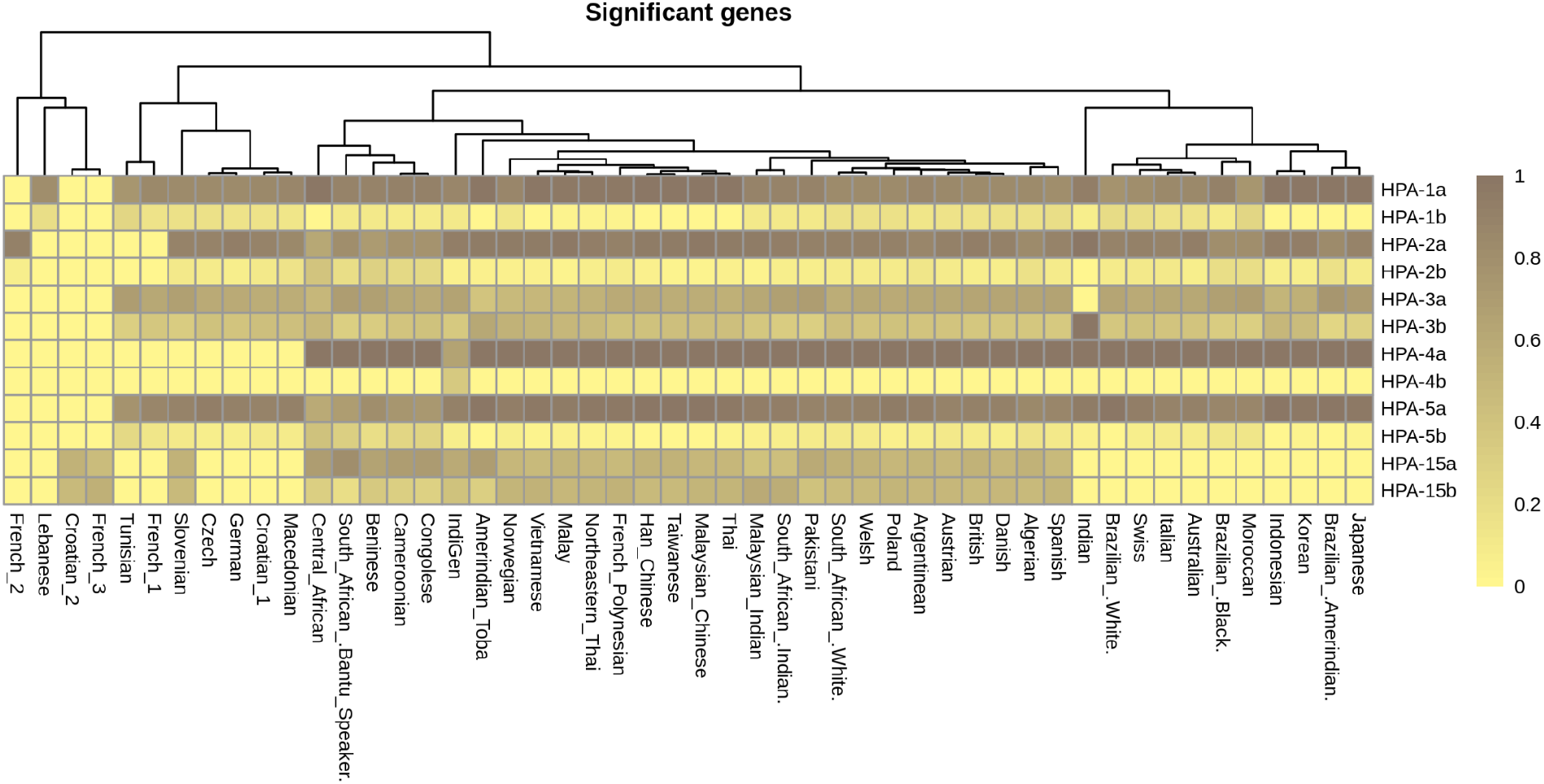
Heatmap illustrating the distribution of HPA allele frequencies among various global populations

## Discussion and Conclusion

Genotyping of human platelet antigens and establishment of population specific donor registries have been suggested to serve in donor selection and management of platelet-specific alloimmunization (Jovanovic Srzentic et al. 2021).

Among the tested HPA systems, HPA-1 and 4 showed significant disequilibrium in the population. Generally, in all HPA systems, “b” allele was observed to occur at much lower frequency in comparison to the “a” allele with the exception of the HPA-15 system where allele frequencies of HPA-15a and 15b were 0.582 and 0.418 respectively. In addition, HPA-4 was the only system with the presence of a “b” allele but absence of “bb” genotype, indicating the heterozygous nature of the allele. Such patterns have been commonly reported in many East-Asian populations (Kupatawintu et al. 2005), (Phuangtham et al. 2017), (Seo et al. 1998), (Asmarinah et al. 2013), (Tan, Lian, and Nadarajan 2012), (Pai et al. 2013).

In comparison to global allele frequencies, HPA-1a and 1b alleles were found similar in most South Asian populations {ref}. Interestingly, HPA-1b allele frequencies varied from very high in populations like Brazilian, Moroccan and Tunisian (0.22-0.25) to low in South East Asians including Malaysians, Thai, Indonesians, Vietnamese, Han Chinese, Taiwanese, Japanese and Koreans. This finding is concordant with previous reports suggesting an increased rate of alloimmunization due to antibodies to the HPA-1a antigen in Caucasians. Allele frequencies of HPA-2a were found to occur in a similar range among all global populations (0.61-0.99) whereas the 2b allele was found at a slightly higher frequency in Africans.

In conclusion, this study provides the first comprehensive catalogue of HPA allele frequencies in the Indian population. Our analysis suggests similarities and differences in the allelic profiles of Indians with that of other global ethinic groups. Such efforts could potentially serve as valuable resources for clinical research and interventions in future.

## Acknowledgements

MI acknowledges fellowship from ICMR-India. MD and BJ acknowledge research fellowships from CSIR-India. The funders had no role in the analysis of data, preparation of the manuscript or decision to publish.

## Supplementary Datasets

**Supplementary Table 1**. Frequencies of HPA alleles observed in various subpopulations of the gnomAD dataset

**Supplementary Table 2**. Frequencies of HPA alleles observed in various subpopulations of 1000 Genomes dataset

## Website references

1. Human Platelet Antigen Database – https://www.versiti.org/medical-professionals/precision-medicine-expertise/platelet-antigen-database

2. IndiGenomes – https://clingen.igib.res.in/indigen/

